# Modeling tumors as species-rich ecological communities

**DOI:** 10.1101/2024.04.22.590504

**Authors:** Guim Aguadé-Gorgorió, Alexander R.A. Anderson, Ricard Solé

## Abstract

Many advanced cancers resist therapeutic intervention. This process is fundamentally related to intra-tumor heterogeneity: multiple cell populations, each with different mutational and phenotypic signatures, coexist within a tumor and its metastatic nodes. Like species in an ecosystem, many cancer cell populations are intertwined in a complex network of ecological interactions. Most mathematical models of tumor ecology, however, cannot account for such phenotypic diversity nor are able to predict its consequences. Here we propose that the Generalized Lotka-Volterra model (GLV), a standard tool to describe complex, species-rich ecological communities, provides a suitable framework to describe the ecology of heterogeneous tumors. We develop a GLV model of tumor growth and discuss how its emerging properties, such as outgrowth and multistability, provide a new understanding of the disease. Additionally, we discuss potential extensions of the model and their application to three active areas of cancer research, namely phenotypic plasticity, the cancer-immune interplay and the resistance of metastatic tumors to treatment. Our work outlines a set of questions and a tentative road map for further research in cancer ecology.

## I. INTRODUCTION

Over the past few decades, a convergence of clinical, experimental, and theoretical cancer research has illuminated the intricate nature of cancer as an ecological, evolutionary, and developmental phenomenon [1–4]. This conceptual framework has provided oncologists with a well-established set of methods and tools from these disciplines, pushing forward our understanding of cancer, its origins and how to better treat it.

The way cancer populations avoid different selection barriers (from physical constraints on the tissue level to immune-related attacks) can be understood as a combination of ecological and evolutionary processes. Much has been written about the latter: well-defined evolutionary events usually punctuate the path towards a malignant tumor. However, the ways these events unfold in time are mediated by ecological change. Once a given barrier is surpassed, new ecological interactions are set in motion and population dynamics determines the outcome of the new context. Ecology is, therefore, at the heart of the many scales relevant to tumorigenesis and cancer treatment, from the growth dynamics of novel phenotypes to the emergence of angiogenic or metastatic properties (Fig. 1) [3, 5–7]. This *cancer ecology* framework has even prompted recent advances in cancer therapy, including the understanding of competitive release and its role in Adaptive Therapy [8] and the unraveling of the complex web of interactions between tumors and the immune system [9].

**FIG. 1:**
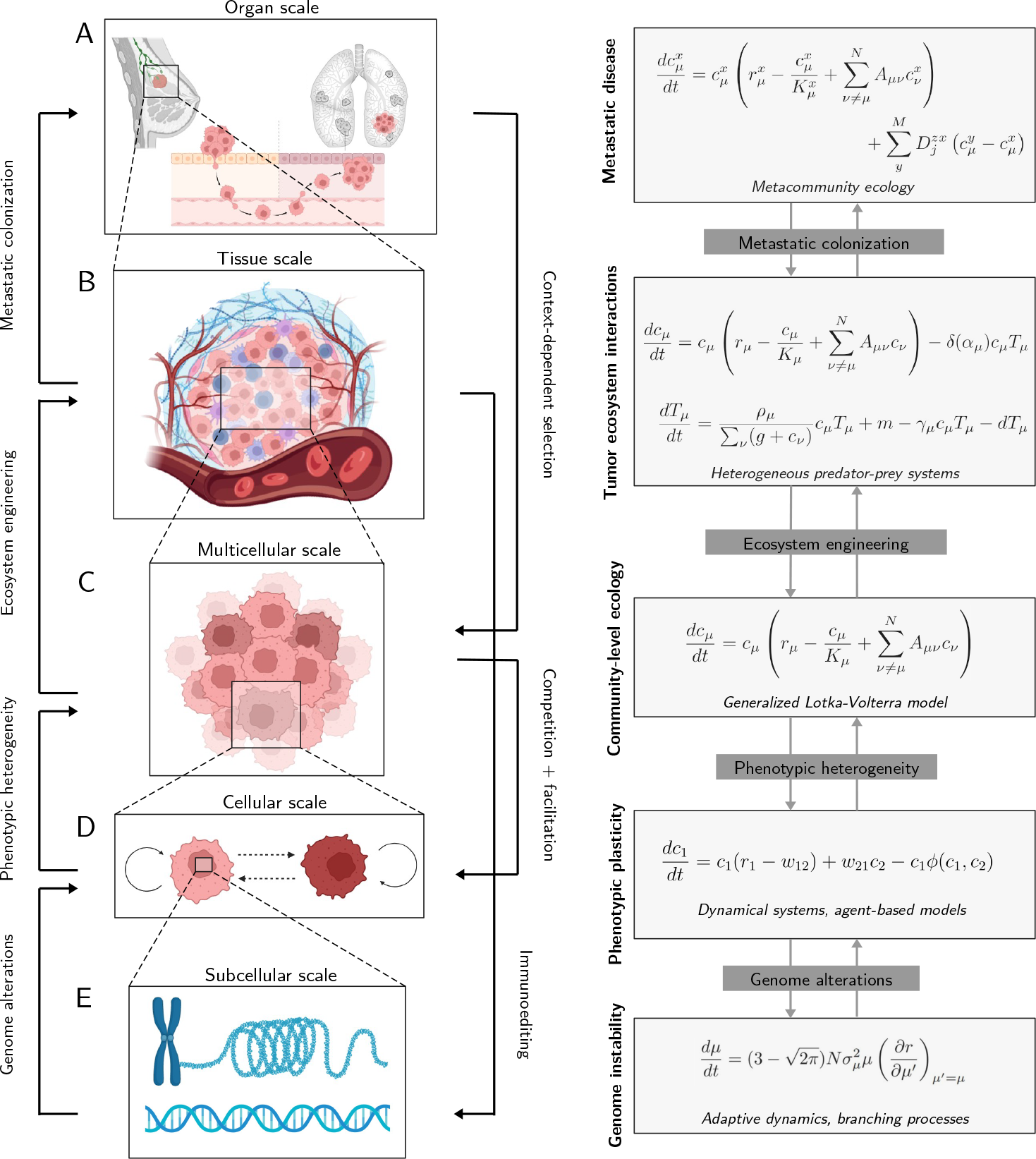
Multiscale complexity, emerging feedbacks and mathematical models of cancer ecology. Multiple scales of complexity participate in the progression towards malignancy, each establishing a set of ecological processes over which evolution operates. We define the layer at play, examples of feedbacks between layers and potential mathematical models able to describe their dynamics. (A) Metastatic colonization at the organ scale [44, 45]; (B) Ecosystem-level interactions with other cell types (in blue) at the tissue scale [43, 46]; (C) Intra-tumor community ecology at the multicellular scale; (D) Phenotypic growth and plasticity at the cellular scale [17, 47, 48] and (E) Genetic, epigenetic and chromosomal alterations at the subcellular scale [49, 50]. While mathematical models in (D,E) are well established, we lack a general framework that can account for the heterogeneity and high-dimensionality of the interaction networks involved in (A-C). Towards this goal, establishing the GLV model as the foundational model of heterogeneous tumors (C) can open the door towards understanding cancer dynamics at higher scales (A,B) [43].

### BOX 1

**Tumor ecosystems as complex adaptive systems**

The ecological nature of cancer cell populations shares a number of universal properties with other complex adaptive systems, and with ecosystems in particular. As far-from-equilibrium, dissipative structures, they exhibit nonlinear dynamical properties that pervade their stability but are also responsible for the structure of the phase space of potential behavioral states. They are in fact *complex adaptive systems* (CAS) [10]. Crucial features that characterize them as CAS would include:

1. Diversity: as it occurs with species diversity in natural ecosystems, intratumor phenotypic diversity is known to play a key role in maintaining robust behavior [11]. While in the former biodiversity acts as a firewall to invader species and is a surrogate of a healthy state, in cancer we understand heterogeneity (tumor diversity) as a source of adaptation [12] and resistance to therapy [13, 14].
2. Non-linearity: as complex ecosystems, tumors exhibit non-linear relationships between components associated to the nature of interactions. Such nonlinear behavior is often characterized by the presence of bifurcation points in parameter space. These special parameter combinations define the boundaries between different *attractor* states, i. e. states (here defined in terms of population abundances) towards which a system tends to evolve over time.
3. Self-organization: tumors have the capacity to self-organize in space and time [15], forming patterns and structures without external control. Self-organization arises from the interactions among individual components, leading to emergent properties at higher levels of organization. These patterns emerge through a combination of both negative (competitive) and positive (cooperative) interactions.

All these concepts have been part of the early development of theoretical ecology [16] and provide the basis to formulate mathematical models that reveal the presence of several key phenomena. These include the stability properties of attractor states and their robustness against perturbations, but also the presence of breakpoints, i. e. parameter values that define sharp changes in the dynamics, as it occurs for example in the transitions to cancer removal under chemotherapy or the activation of immune responses against neoantigens.

Mathematical models of cancer ecology provide us with an explanatory framework that offers both qualitative understanding of the underlying mechanisms as well as potential predictive power. By exposing the logic of cancer population dynamics, they can help explain how and why therapies succeed or fail. These models have been traditionally built as coarse-grained descriptions of cell populations, often reducing them to more or less homogeneous compartments. Their simplicity allows for a full understanding of their implications, providing a descriptive link between the ecological rules at play and the growth of the tumor [3, 17–20] (Fig. 1D). In some cases, the tumor is represented as a single population of cancer cells [17] while in most studies the interactions between cancer cells and other cell populations (from healthy tissue to the immune system), two-dimensional models have successfully illuminated crucial aspects of the nonlinearities involved [19, 21–23]. However, the improvement in our understanding of the richness of tumor diversity has led to the realization that cancer population dynamics makes sense under a *community* ecology picture [24]. In this context, heterogeneity at the mutational, phenotypic and cell type levels, dictates that tumors are in fact complex adaptive ecosystems built of many interacting populations (See BOX 1, [25–27]). Furthermore, despite the traditional dominance of competition as a driver of cancer dynamics, evidence accumulates indicating that ecological interactions between cancer cells are not only competitive: cooperation or commensalism could also be at play [25, 28–30]. More importantly, the combination of dedicated experimental efforts and improved mathematical models has revealed a multiscale picture of cancer where novel phenomena emerge as we move across scales (Fig. 1).

Mathematical oncology has made a great progress over the last decades, where a whole array of theoretical approaches has been developed [3, 31–34]. Along with population dynamics in space and time, approaches applying evolutionary game theory have also been helpful in understanding the emergence and interplay of different cancer phenotypes [35–38]. With the rapid development of -omics data and improvements in single-cell level analysis, the distance between theory and experimental and clinical information has been rapidly shrinking. However, there is no current consensus on a mathematical framework that upscales simple models towards describing the species-rich cancer ecosystem (Fig. 1A-C). Following its widely accepted application on ecological and microbial systems [39–42], we propose that the high-dimensional model defined by the so-called Generalized Lotka-Volterra interactions (GLV) is a powerful candidate to understand and predict the complexity of tumors as ecological communities [43].

In this paper we present the emerging properties of the GLV model and their implications in tumor population dynamics. We first describe the model itself and its application in cancer ecology (Section II.A). In particular, the model captures the conditions for phenotypic coexistence and tumor outgrowth (Section II.B), as well as the possibility of community-level succession and shifts between multiple cancerous states, and new questions regarding treatment administration (Section II.C). Second, we propose three additional applications of the species-rich GLV cancer model on high-dimensional phenotypic plasticity (Section III.A), cancer-immune interactions (Section III.B) and the metastatic process (Section III.C).

Inspired by the pioneering work of Robert Gatenby on the application of ecological models to cancer research, *Population ecology issues in tumor growth* [17] (Fig. 1D), our work can be read as an updated proposal of *Community ecology issues in tumor growth* that takes into account our current understanding of cancer complexity (Fig. 1A-C).

## II. THE MATHEMATICS OF HETEROGENEOUS TUMORS

Due to their diverse phenotypic makeup, heterogeneous tumors can display intricate dynamic patterns over both space and time. This phenomenon has been a focal point in ecology, with a longstanding tradition addressing this challenge [51]. Particularly, the use of multispecies models based on mass-reaction kinetics, as encapsulated by the GLV equations, has been prevalent [39, 52–54]. These mathematical frameworks have been instrumental in comprehending the dynamics of real ecosystems, aiding in the elucidation of underlying organizational principles. They have played a pivotal role in making sense of counter-intuitive findings, such as the relationship between diversity and connectivity [55], and in understanding how ecosystem resilience and species composition evolve with gradual changes in environmental variables [56].

### A. The generalized Lotka-Volterra model of cancer growth

The GLV model is a benchmark description for the dynamics of ecological communities of many interacting species [57]. Here we present the main features of this model and its application to cancerous tissue. The model captures the abundance of each of *S* species as a function of its adaptation to an abiotic environment and its interactions with the rest of the species in the community, as weighted by the so called communiy matrix 𝒜, defined as:

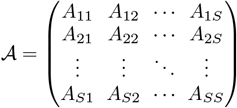

The abundance *N*_*i*_ of each species then follows the set of *S* coupled nonlinear equations:

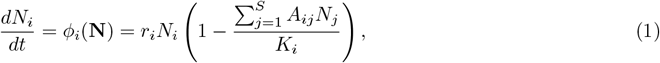

where *r*_*i*_ is the intrinsic rate of replication, while *A*_*ij*_ encodes the possible effects of species abundances **N** = (*N*_1_, …, *N*_*s*_) on the growth of *N*_*i*_ [39]. Both *r*_*i*_ and the diagonal elements *A*_*ii*_ capture the adaptation of a species to its niche, describing its dynamics when alone, while *A*_*ij*_ captures the biotic interactions between different species. This model contains several limit cases that describe standard, low-dimensional scenarios (see BOX 2). The relevant attractor states *S*^*^ (possible stable community configurations) are obtained from the condition *dN*_*i*_*/dt* = 0 (for *i* = 1, …, *S*, which provides a set of vectors:

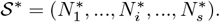

Determining the set of attractors and their stability properties (see [53, 58] for a standard treatment and methods) will be at the core of all the models presented here.

In many ecological communities, the number of species *S* and the complexity of their interactions make it extremely hard to empirically obtain all parameters of Equation (1). Instead, mathematical models of species-rich ecosystems have proposed that interactions are in fact so complex that details cease to matter and one can ask whether generic patterns emerge [40]. Since the early work of Robert May, this has been studied by modeling the elements of *A*_*ij*_ as randomly distributed variables [55], a method that has brought fundamental insight into the functioning and stability of complex ecosystems (see e.g. [40, 57]). Furthermore, the last decade has seen an outburst of analysis for this apparently simple but high-dimensional model (see e.g. [39, 40, 59–61]). This provides cancer researchers with a rich framework to understand what can happen when many cells interact under competition, cooperation or predator-prey mechanisms.

How does the model translate when trying to describe a tumor population? In low dimensional form it has been thoroughly applied to the study of tumor growth and competition (see Box 2, [17, 62]). We propose here that each cancerous population, defined by a set of signatures yielding a functional phenotypic behavior *μ* [12, 27, 29, 43], needs to be understood as an individual species with abundance *c*_*μ*_ growing while interacting with the other cellular populations *c*_*ν*_. Depending on the dynamics at play, here *c*_*μ*_ could be restricted to populations with equivalent mutational or antigenic features (so-called genetic clones, [12]), or even include additional layers of the non-cancerous tissue such as the stroma [63] or the immune system [43]. A minimal expression for these dynamics can be written as

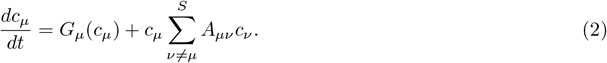

#### BOX 2

**Competitive dynamics between two populations**

The general equations introduced by the GLV model (1) contain all the classical, low-dimensional examples of ecological dynamics, including single-species growth, two-species competition or two-species mutualism. For *S* = 1, using *A*_11_ = −1 we obtain the logistic equation (which is one of the possible candidates for cancer growth, see [64–66]), whereas for *S* = 2, using the matrix

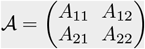

and assuming *A*_*ij*_ *<* 0 we obtain the standard Lotka-Volterra two-species competition, which could represent the competition between cancer and healthy cells [67, 68]. One particularly simple case is provided by the symmetric scenario, where *r*_*i*_ = *r*, self-regulation occurs with strength *A*_11_ = *A*_22_ = −1 and *A*_12_ = *A*_21_ = *β <* 0. For this particular case, the model already exhibits four equilibrium (fixed) points obtained from the condition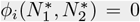. These points define four possible *attractor* states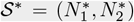, namely: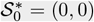 (extinction), 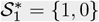 and 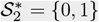(exclusion points) and 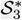 coexistence state where both species are present at 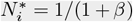. If we analyse the stability of these points, it is found that, for *β <* −1, the only possible stable solutions are the exclusion points 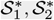: competition is so strong that the population that starts at higher abundance will outcompete the other. The coexistence alternative is obtained when *β >* −1. In that case, it can be shown that the system exhibits stable coexistence and the exclusion points are unstable. For *β >* 1 we have outgrowth (and the system diverges). Other kinds of pairwise clonal interactions in cancer can be defined, including comensalism and even parasitism. Their relevance for quantitative clinical studies and their connection with different mathematical approximations in summarized in [25].

The dynamics of tumor cell populations is split into a growth term *G*_*μ*_ on the one hand and the interactions with the rest of populations on the other. Consistent with the view of sigmoid growth saturating at high abundances [62, 69], a reasonable assumption is a logistic growth curve of the form *G*(*c*) = *c*(*r* − *c/K*) [62]. Both *r* and *K* are likely to be heterogeneous variables, as different phenotypes can modulate their rate of replication and the available nutrients or density constraints of their environment [70]. This mapping between logistic growth in tumors and the original GLV model will be followed throughout the rest of this paper. However, it is interesting to note that other saturation functions have been tested against tumor growth data, with the Gompertz law and sublinear allometric scaling being prominent examples [62]. In this context, while exponential growth has been found for example in leukemias and lymphomas, If surface effects become relevant the growth pattern departs from the exponential shape. When proliferation occurs in a system where spatial effects (such as surface ruggedness) is at work, *G*(*c*) should take the form

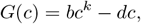

with *k <* 1 [62, 71]. Such sublinear growth terms are known to affect the resulting species diversity, as it has been explored within the context of replicator dynamics [71–73].

Beyond population-level growth, the GLV model explicitly accounts for heterogeneity and interactions at the phenotypic level. The simplest possible description is that each cancer population interacts with the rest of the tumor through a complex network of inter-species linear growth effects encoded in *A* (Fig. 2A). Beyond the well-accepted view of cancer clones competing for resources and space (*A*_*μν*_ *<* 0) [25, 70, 74], recent research highlights that tumors also harbor cooperative interactions (*A*_*μν*_ *>* 0), resulting from the secretion of shared growth factors and inflammatory signaling[25, 28, 43, 75], as well as commensalism, where other populations freely benefit from an angiogenic, invasive or metastatic phenotype (*A*_*μν*_ *>* 0, *A*_*νμ*_ = 0) (Fig. 2A) [25, 76].

**FIG. 2:**
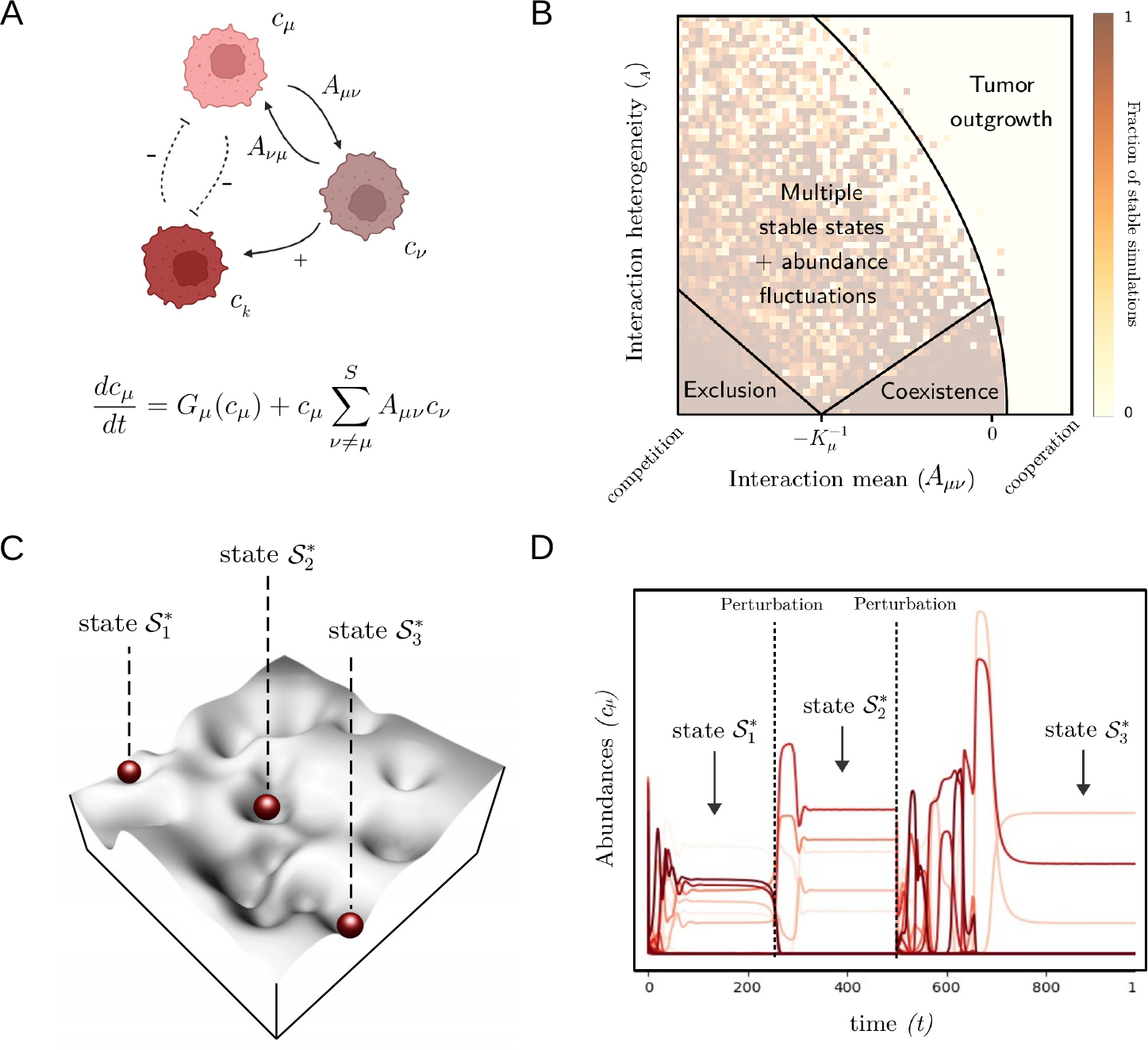
Tumors as species-rich ecological communities. (A) Many phenotypic populations *{c*_*μ*_*}* coexist within a tumor and interact under multiple ecological processes encoded in *{A*_*μν*_ *}*. The GLV model (lower formula) is a suitable model to describe such high-dimensional tumor communities. A crucial concept in this modeling approach is the presence of distinct dynamical regimes (B), each emerging for given mean interaction strength *(A*_*μν*_ ) and interaction heterogeneity *σ*_*A*_ values. The model harbors both a phase of unique stable coexistence, as well as a particularly relevant regime of community multistability, where multiple attractor states can coexist, as represented in (C) with the minima of the landscape associated to multiple stable states. In (D), we simulate the trajectories of 50 interacting populations in this multistability regime and apply two perturbations that decrease the abundance of all populations at random (dashed lines). Even if each perturbation strictly reduces the number of present cells (e.g. a treatment shock), community dynamics can lead to shifts from an heterogeneous tumor community 𝒮_1_ towards states with higher abundances 𝒮_2_ and 𝒮_3_.

Although initial endeavors are underway to measure the interaction strengths encoded in *A* within tumors [25], accurately mapping intercellular effects *in vivo* remains a formidable challenge. Given the intricacies of tumor complexity, the GLV modeling strategy, which assumes for a probabilistic distribution of random *A*_*μν*_ values, emerges as a promising approach. This strategy has already proven successful in describing microbial communities [42]. By applying this methodology to cancer experiments, we could gain fundamental insights into intra-tumor dynamics solely by approximating the statistics of *A* (see Fig. 2B, [25, 77, 78]).

The use of the variance of interactions as a key parameter to study the potential dynamical states of ecological networks has been recently incorporated. What are the outcomes of this high-dimensional view of cancer? A key feature of the GLV model is that different dynamical phases emerge depending on the mean interaction strength, i. e.

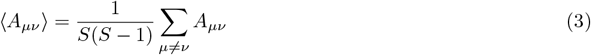

and the standard deviation of the values in *A*, namely 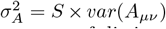. These two statistical average values allow to define a partition of the (⟨*A*_*μν*_⟩, *σ*_*A*_) plane into a set of distinct phases. Below we describe how the emerging dynamical phases can have direct implications for our understanding of cancer.

### B. Coexistence, outgrowth and competitive release

Four distinct dynamical phases can be discerned within the GLV model, as depicted in Fig. 2B [39, 61]. These phases encompass stable species coexistence, exponential growth leading to competitive exclusion, described here, and a more intricate phase marked by fluctuations and multistability, to be described further below. Each phase illuminates a distinct aspect of the cancer development process.

Understanding how ecological communities can stably maintain a high number of coexisting species has troubled ecologists for decades [79–81], since theoretical predictions would expect competitors to exclude each other [82], and multiple consumers require as many different resources to survive [83], or species-rich communities to become unstable at high diversity [55]. This question is also relevant – and almost equivalent – in cancer research [84, 85]. Many phenotypic and mutational profiles coexist inside tumors [12, 86, 87], while we would expect selection to allow for only a few [88].

The first phase of the GLV model (Fig. 2B) is that of *stable coexistence*: when interspecies competition is not as strong as carrying-capacity regulation, i. e.

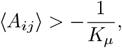

(Fig. 2B), a large fraction of cancerous phenotypes is able to coexist [39, 55]. This characterizes an early phase of tumor growth, where physical or nutrient constraints (and hence low carrying capacity and strong self-regulation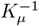) keep pre-cancerous populations under a homeostatic-like equilibrium [85, 89]. Moreover, consistently with observations in ecological communities, heterogeneous species-abundance distributions are observable in this phase. indicate that a few species are extremely abundant while most are rare [90]. This brings forward a potential prediction of the GLV formalism for the abundance distributions of cancer cell phenotypes.

Once evolution kicks in and self-regulation *K*_*μ*_ is no longer the limiting factor [70], the GLV predicts the emergence of richer dynamical phases. In the context of extreme intra-tumor competition driven by large fitness differences, the system transitions to a regime of *competitive exclusion*, where communities are dominated by a single, strongly competing phenotype (Fig.2B). This regime is particularly relevant to treatment design. The framework of Adaptive Therapy (AT), for instance, has shown how maximum dosage treatment, by eradicating drug sensitive populations, provides a competitive advantage to the drug resistant population [17, 20]. AT instead aims at adapting drug dosage and tempos to maintain the tumor within the controlled coexistence phase (Fig.2B). Given that multiple phenotypes might be grouped into the sensitive compartment, we hypothesize that the multispecies GLV model better predicts the conditions by which a mixture of sensitive populations can keep the resistant phenotype at bay [91].

The third well-known phase of the GLV model considers *tumor outgrowth*: under heterogeneous interactions involving weak competition but also a certain degree of cooperation, a fraction of phenotypes will expand exponentially, as empirically observed in [29]. This regime likely explains early to intermediate stages of tumor growth, where abundant resources and weak spatial competition coexist with cooperative signatures [29, 75]. Consistent with recent findings [29], the GLV model provides an explanation for how, even within stressed and competitive tumors, high heterogeneity can drive the emergence of cooperation leading to sudden tumor outgrowth after a long period of apparent dormancy [92]. This transition would emerge when a new phenotypic signature displaces community interactions *A* towards crossing the coexistence-to-outgrowth threshold, through e.g. angiogenesis [28], immune evasion [93] or a shared invasive phenotype [29] (Fig. 2B).

### C. Succession and shifts between multiple cancerous states

In many tumors, intraspecies and interaction parameters across phenotypic profiles are likely to be highly heterogeneous [12, 94]. Moreover, in advanced stages of resource-limited tumors, competition is also expected to be strong: persistent acifidication of the environment, reduced resources or increased crowding likely induces negative *A*_*μν*_ values [95].

In this context of competition and heterogeneity, the GLV model opens the possibility for a new paradigm in cancer ecology. In the model, heterogeneity can drive the system towards a complex phase of phenotypic fluctuations and shifts between multiple community states ([39, 60], Fig. 2B), as recently tested in experimental microbial communities [42, 96]. What are the implications of these emerging dynamics for cancer research?

Opposite to the notion of homeostatic stability of a single precancerous state, here heterogeneity drives the system to a much more nuanced scenario. In it, evolutionary changes [12, 97] or persistent phenotypic switching [98] would drive the system towards a metastable regime, in which the whole tumor community can transition through a set of multiple cancerous states [99]. In ecology, this is linked to the concept of ecological succession, by which ecosystems can follow directional – albeit not always predictable [100]– transitions from one ecosystem state to the next [99, 101]. In cancer, recent research highlights the possibility that tumor ecosystems explore recurrent and potentially step-wise predictable trajectories in their composition and malignancy [4]. In this context, the GLV model can provide a first mathematical framework explaining how successive oncogenetic stages and the shifts between them emerge in heterogeneous cancers. A powerful framework could result from connecting cancer models with the structural stability approach [102], which has been successfully applied to explain switching patterns in microbiome dynamics [103].

Beyond explaining the non-trivial natural histories of tumors [4, 104], the possibility of multistationarity in cancer also holds key implications for therapy. In this phase, the extinction of a targeted population under treatment could induce secondary extinctions and invasions [105], yielding an abrupt shift towards a different phenotypic community. In ecology, this is characteristic to the notion of *communities as superorganisms*, where the survival of each species is strongly intertwined to the presence or absence of other species [106]. If tumors behave as GLV superorganisms, this would mean that a whole new ecological mechanism of community-level plasticity is at play without the need for mutational or epigenetic alterations. Abrupt transitions from one phenotypic composition to another imply that not single phenotypes [20], but entirely new resistant communities could emerge after a failed treatment attempt.

Overall, the possibility that heterogeneity drives tumors towards a non-trivial phase involving whole-community shifts urgently asks for a more dynamical understanding of cancer and its complexity if we are to design successful therapeutic strategies [107].

## III. ONCOLOGICAL EXTENSIONS OF THE GLV MODEL

### A. High-dimensional phenotypic plasticity

The parallel between species-rich communities and heterogeneous tumors suggests new, unexpected cancer properties. Yet, there are characteristics of tumor cells and their plastic genome that cannot be described by classical ecological dynamics. For example, accumulated evidence indicates that rogue cells do not necessarily express a stable phenotype (as animal or plant species do), but are much more plastic and can stochastically switch into others [98, 108]. A relevant example here is the recent observation of a complex architecture of four well-defined switching phenotypes in Glioblastoma [109]. How can we mathematically characterize such a system, and what are the implications for therapy?

Inspired in minimal models of bacterial plasticity [110], now each population *c*_*μ*_ switches at rate *W*_*νμ*_ towards other *c*_*ν*_ phenotypes, and receives new individuals from them at rate *W*_*μν*_ (Fig. 3A). The diagonal elements of the *W* matrix are zeros, as self-switching does not exist, and off-diagonal terms are not necessarily symmetric [98]. The previous dynamical equation for the interaction and growth of tumor populations now reads:

**FIG. 3:**
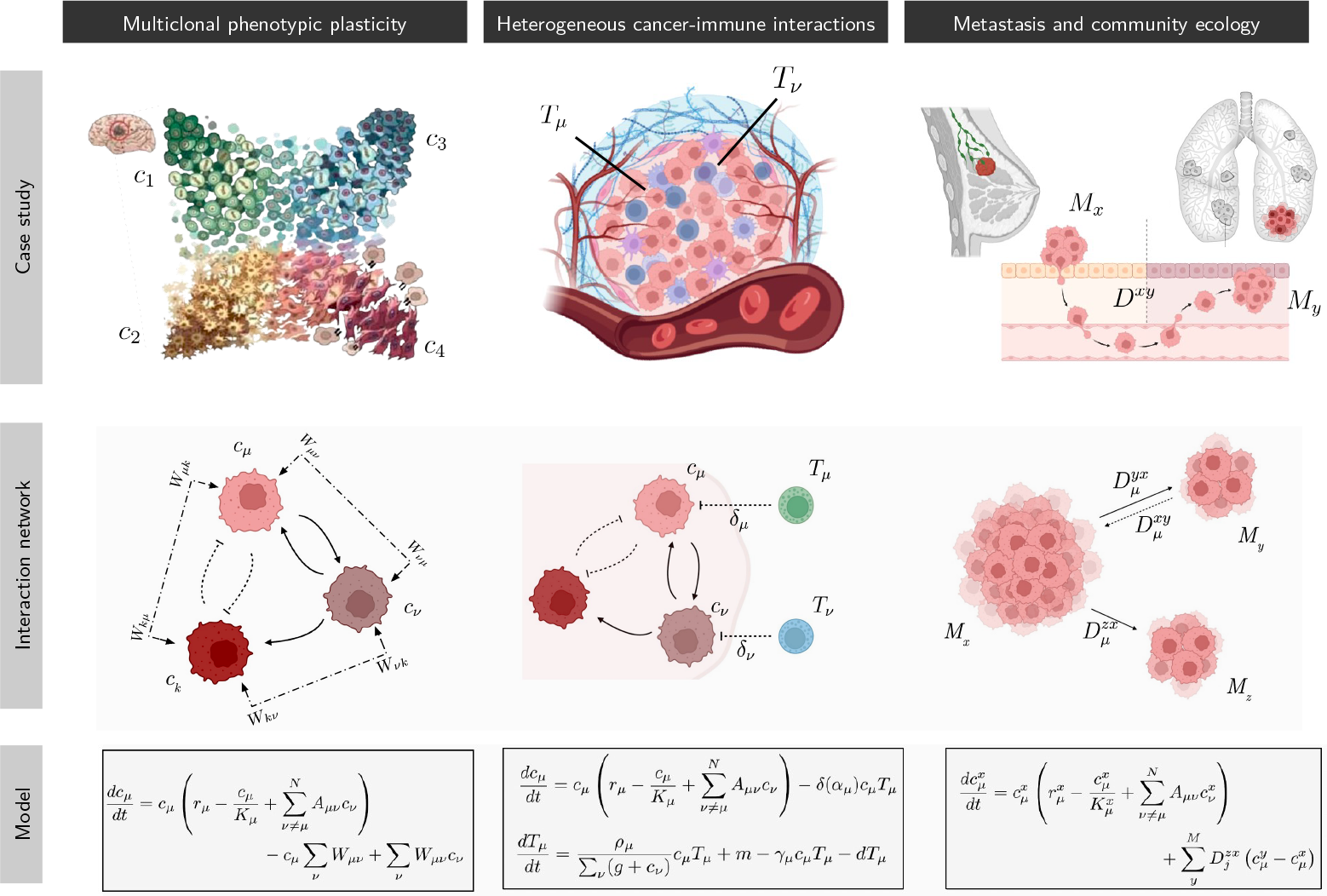
Oncological extensions of the GLV cancer model. The species-rich nature of tumor communities holds implications in other areas of cancer research, as exemplified here with three case studies. First column: Ecological interactions between cellular phenotypes are sometimes intertwined with stochastic phenotypic plasticity, as it occurs with glioblastomas, where genetic analysis revealed four transitioning phenotypes [109]. These transitions can be described by a matrix of rate flows between population compartments indicated by the terms *W*_*μν*_. Second column: Interactions between cancer and the immune system are inherently high-dimensional, and complex patterns of cancer-immune interactions appear to be a common feature. Multiple cancer clones *{c*_*μ*_*}* interact with corresponding T cells *{T*_*μ*_*}*, where each T cell species can only recognize and attack one immunogenic antigen. Third column: The problem of metastasis can be modeled as a metacommunity with multiple patches *{M*_*i*_*}* connected by cell dispersal 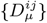, yielding emerging properties of the tumor-nodes system.

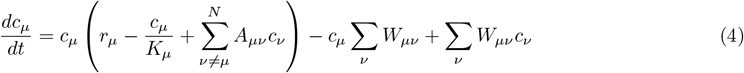

A first approach to understand the added complexity is to consider minimal scenarios with two phenotypes at play, such as the study of epithelial-mesenchymal plasticity and its implications in metastatic spreading [111] or the analysis of a drug sensitive-drug resistant switch [112, 113]. Low-dimensional models involving a two-phenotype switch following equation (3) predict that therapy can only succeed if it targets the switching matrix *W* [114]. Treatment aiming at cell division or death alone will fail if phenotypic switching is at work, as this will maintain the heterogeneous community in place [47, 115]. As seen for bet-hedging strategies in bacteria [110] and closely connected to game theory models [38, 41, 115], phenotypic switching models predictably explain how plasticity emerges as a community-level resistance strategy.

What happens when more than two populations can actively switch phenoytpes [109, 116]? Recent analysis of combination therapy models following a simplified form of equation (3) provide a prediction for which treatment scheme is most effective [113, 115]. In particular, the species-rich model predicts not only the necessary targeting of switching rates, but also a threshold number of drug sensitive phenotypes below which the tumor can escape treatment [115].

Even if preliminary results already indicate the potential of phenotypic plasticity as a resistance mechanism, the complete picture painted by Eq. 3 is yet to be fully understood. When both ecological interactions (*A*) and stochastic switching (*W*) are taken into account, persistent phenotypic switching between populations might not only prompt resistance, but also community-level shifts towards alternative tumor architectures where one or another phenotype dominate the system [109, 116]. Theoretical analysis of the full system (3) should search for potential weak spots to this otherwise robust and plastic system: treatments for phenotypically plastic tumors are likely to fail unless such emerging complexity is fully accounted for.

### B. Heterogeneity in cancer-immune interactions

Uncovering the ecological interactions between cancer cells and the immune system has prompted revolutionary changes in cancer therapeutics. The immune system, in fact, is itself an ecosystem of cellular species interacting in complex ways to mediate the immune response [117]. In cancer, different immune cells participate in tumor cell recognition and destruction [18, 118], but also in pro-tumor inflammatory responses [7].

One of the most relevant components for cancer treatment are adaptive T cells and their complex recognize- and-kill cascade (Fig. 3B) [118]. Early ecological models proposed that cancer-T cell interactions behave as a predator-prey system [18, 22]. Also fundamental to this process is the discovery that cancerous mutations, through the alteration of surface antigens, can activate T cell recognition [119]. This poses a selective barrier, where tumors that escape the immune system are more likely to avoid early deletion [93, 120].

Accumulated evidence indicates that one escape mechanism is rooted in tumor heterogeneity: it is not only the number of neoantigenic mutations, but also their variety what dictates the failure of the immune response [121, 122]. Here we propose extending the GLV cancer model to account for immune attack. One possible starting point is the two-dimensional cancer-immune model where both predation and competition are in place [21, 22], described by the dynamical equations

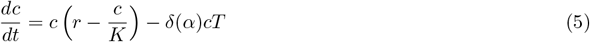

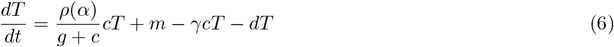

where the last term in *c*? describes death by immune recognition and attack at rate *δ*(*α*) related to the neoantigen load of the tumor *α* [46]. T cell growth due to neoantigen recognition increases as *ρcT* (a larger bulk *c* will be more easily detected), but only up to a certain value: larger tumors will be harder to penetrate by the immune system [22]. Recent evidence, however, indicates that ecological interactions between different cell types at the tissue scale are also pervaded by heterogeneity (Fig. 1B). In the context of lymphocitic interactions, the response of tumors to immunotherapy is strongly related to how homogeneous their neoantigen load is [46, 121]. The GLV model provides a methodological way to account for neoantigen heterogeneity in cancer. If each cancer population *c*_*μ*_ is now defined by its mutational subclonal profile, yielding a specific neoantigenic component *α*_*μ*_, and *T*_*μ*_ is each T cell population recognizing the most immunogenic neoantigen in *c*_*μ*_, the dynamics of the system can be reduced to (see [43, 46]):

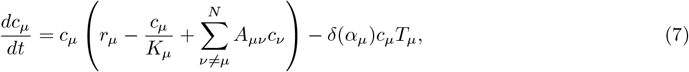

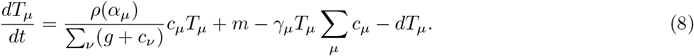

Now tumor populations interact through the GLV model with an additional predatory component (Fig. 3B). However, it is key to observe that the predation-competition scenario is not even: cancer populations *c*_*μ*_ are predated by the immune compartment that can recognize them *T*_*μ*_ [122], whereas T cells die in the presence of whatever cancer populations are in place, Σ _*μ*_*c*_*μ*_. Furthermore, tumor clones avoid external attack by cooperatively increasing the tumor bulk, Σ_*ν*_ (*g* + *c*_*ν*_) [123]. Each T-cell clone recognizes one cancer neoantigen, while all cancer clones can kill T cells and also cooperate to avoid immune assault [46]. Is there a limit beyond which such a *divide and win* strategy succeeds? Can it explain why immunotherapy is so dependent on neoantigen load and immunogeneicity?

Early models of HIV progression uncovered a similar problem: there is a critical viral diversity beyond which the immune system can no longer control viral growth [124]. Similarly, a recent attempt at tackling the complexity of equations (6-7) indicates that the extended GLV model contains a neoantigen heterogeneity threshold [46]: if the number of different neoantigenic clones overcomes this critical value, the tumor will become too heterogeneous to be effectively targeted by the immune system [121, 122]. When applied to clinical data of melanomas treated with immunotherapy, this threshold value consistently predicts which patients will respond better to immune blockade treatment [46].

The whole network of cancer-immune interactions, however, spans much beyond the predator role of T cells. T cells themselves can also hold tumor promoting phenotypes, yielding non-trivial scenarios where tumor cells become engineers of their own microenvironment [7, 43]. Similarly, the role of macrophages in cancer is inherently multimodal: type-1 macrophages interact in a cooperative antiinflammatory cascade with tumor cells, while type-2 macrophages can eradicate rogue cells [7]. Beyond immune cells, stromal recruitment is another of the many layers that participate in dynamics of the tumor ecosystem ([125], Fig. 1B). Cancer vaccines, which modulate the landscape of immune recognition of neoantigens, could also be introduced [126]. As done for neoantigen heterogeneity biomarkers in melanoma [46], including these additional layers in the complex ecology of Eqs. (6-7) could bring new insight into the tumor ecosystem network and how to modulate it to treat cancer.

### C. Metastasis and metacommunity ecology

Advanced stage metastatic disease accounts for the majority of cancer-related deaths [91]. On top of the aforementioned cellular heterogeneity, seeding between the primary tumor and multiple metastases colonizing different organs imply an ever more complex disease that is inherently difficult to understand and treat [127] (Fig. 1A). In this direction, mathematical and ecological theory have shed light into different aspects of the process [44, 127], such as elucidating how phenotypic differences between the primary tumor and its metastatic nodes can inform of the seeding mechanisms at play [128]. Ecologically, the complexity of the problem is that we no longer have a community adapted to a given niche (the microenvironment of the host organ, Fig. 1B), but rather a set of heterogeneous tumor communities connected by cell migration [91] and adapted to alternative microenvironments [92].

Since the 1990’s, ecologists know that a *metapopulation*, a population distributed along spatial patches connected by dispersal, can display dynamics not found in single-patch systems [129]. In particular, while stochastic birth-death processes can drive single populations to extinction, enough migration between patches allows the metapopulation to thrive, yielding a so-called *rescue effect*. Early work already indicated that metapopulation dynamics could be at play in heterogeneous tumors and provide an explanation (based on spatial ecology) for the coexistence of diverse clonal cancer cell populations [86]. In the context of metastatic disease, this emerging property could explain how cells arriving from the primary tumor, or even seeding between metastatic nodes [91], could allow weaker or targeted metastases to survive under therapy.

What happens when host-level disease is not a single ecological community (Fig. 1C), but many communities connected by migration (Fig. 1A)? Will the metastases replicate the phenotypic composition –and hence the treatment sensitivity– of the main tumor, or else can theory help explain if each metastasis forms a community of its own? We propose here that the complexity the metastatic process can be fundamentally captured by the theory of metacommunity ecology (Fig. 3C). Metacommunity ecology is an extension of metapopulation ecology that studies a network of geographic patches connected by species migration, where each patch is itself a community of interacting species [45]. The GLV system is extended to a network of *M* communities (the primary tumor and metastatic nodes), where the abundance of phenotype *μ* on node *x*, 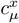, depends on its GLV dynamics but also on the dispersal from and towards the rest of the nodes (Fig. 3B, [130]):

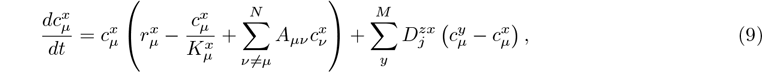

where 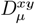 captures the migration of *μ*-cells between the nodes *x* and *y*. Current research allows us to accurately capture how the tumor seeds the different metastatic nodes [128], providing a glimpse on this key property of the metacommunity. Here we highlight three recent results on species-rich metacommunity ecology that could provide novel predictive tools to understand metastatic cancers once *D* can be estimated.

First, and inspired by island biogeography [59], the GLV model with cellular seeding from a large phenotypic pool could be seen to describe a single metastatic community seeded by the main tumor. The model allows us to establish a link between the phenotypic composition of the metastases and the primary tumor, and how both are impacted by species interactions and the tissue microenvironment at each site [59].

However, measuring the interactions and environmental impacts *in vivo* is a very difficult task. In this context, recent results show that species distribution patterns (in cancer, how different phenotypes are distributed along the primary tumor and its metastatic sites) can already predict certain properties of the niche differences and the community interactions at play [130]. Applying these results to cancer data could help unravel if phenotypic distributions, which sometimes show metastases that are very similar or different from the original tumor [128, 131], can inform about the microenvironment at each metastases. This, in turn, could help design combination therapies able to target the heterogeneous adaptation strategies that have emerged at each site.

Third, recent results are elucidating how the rescue effect upscales when multiple interacting populations are at play. The model shows that species migrations, combined with the heterogeneity of their interactions, could allow the metacommunity to survive even further than a metapopulation would, thanks to the positive effect of interspecies cooperation [132]. When translated to oncology, multicellular rescue effects between metastatic compartments could provide key insight into how and why heterogeneous metastatic disease is so difficult to treat. Again, species-rich ecology needs to be considered if we are to design successful therapies.

## IV. DISCUSSION AND OPEN QUESTIONS

Ecological interactions shape all stages and scales of tumor progression (Fig. 1). Beyond the usual focus on the cancer cell, evidence indicates that tumors are built upon a rich and heterogeneous set of populations interacting under ecological mechanisms. In this context, current one-or few-species ecological models of tumor growth cannot capture the complex dynamics of cancer progression.

We propose to upscale current ecological models of tumor growth by applying the mathematical theory of species-rich ecological communities. Our central hypothesis is that the GLV model and its variations provide a candidate framework to describe cancer dynamics, and that its emerging properties shed new light into different phases of tumor progression.

The GLV model provides an advanced toolset to predict the conditions that dictate multispecies coexistence, disease outgrowth and competitive release after treatment. Moreover, the model harbors a non-trivial phase, where heterogeneous interactions can drive the tumor ecosystem towards shifting between multiple cancerous states. If cancer does behave as a GLV community, these salient features could provide a fundamentally novel view on cancer as a plastic and persistently changing complex ecosystem. In this perspective, we propose that modeling tumors as species-rich GLV communities opens the following novel research avenues and questions in mathematical oncology:

1. The diversity-stability debate in ecology plays a key role when modeling pervasive tumor heterogeneity. Given that cancer populations might grow following sublinear dynamics, can this explain and predict the degree of heterogeneity a tumor can support?
2. Species-rich models, when confronted to simpler dynamics, uncover the emergence of multiple stable states. Given that cancers are ecosystems of many interacting species, could transitions between multiple cancerous states provide an additional mechanism for tumor resilience?
3. Tumor phenotypes might be much more plastic than ecological species. What are the impacts of this additional layer of complexity in making cancers difficult to treat?
4. Tumor-immune competition is strongly governed by the neoantigen heterogeneity of the cancer bulk. Can there be predictable thresholds, beyond which the predatory role of T cells is impaired by excessive antigenic diversity?
5. Metastases progress by establishing novel cancer communities in different microenvironments. Can meta-community ecology, the framework to describe ecosystems connected by dispersal, explain the heterogeneity and resilience of the tumor-metastases system? Despite our work being focused on the above topics, the GLV formalism offers multiple paths for further exploration and many relevant open questions regarding diversity dynamics, the role of space and time, the presence and implications of critical points or the origins of plasticity and robustness of tumors. Some relevant possibilities not discussed here are:
6. The models presented here share a deterministic character, but stochastic dynamics can be implemented by generalizing the previous equations [133]. One way is to write down the GLV competition scenario as follows: [39, 134]:

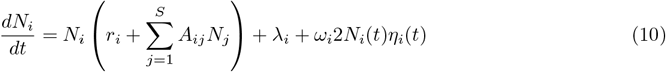

where the last term on the right hand side introduces stochasticity as demographic noise [135]. The term *λ*_*i*_ stands for immigration in an ecological context of a system that is connected with some geographical species pool, but could also be used to introduce mutational events. A relevant result in this context is that, when species diversity increases, competitive communities have an landscape pervaded by marginal attractor states leading to complex fluctuations [134] (see also [136]). Could cancer cell populations evolve towards these marginal states?
7. The attractor landscape that we have presented here can be challenged by the presence of long transient phenomena [137]. It has been known for a long time that the convergence to a given attractor state can be strongly affected by the nature of the nonlinearities, stochastic fluctuations as well as spatial degrees of freedom [138, 139]. The role played by space and long transients has been shown to provide opportunities of ecosystem management [140]. Could these phenomena play a role in heterogeneous tumors?
8. Spatial interactions have been shown to introduce novel properties in the dynamics of complex ecosystems. A whole research area within theoretical ecology is devoted to *spatial ecology* [141, 142]. The presence of spatially explicit metapopulations provides a dramatic example of how space modifies the expectations from well-mixed (mean field) approximations. An example is competitive exclusion: under the presence of space, this process becomes local and global coexistence is possible [143]. Similarly, the locally constrained nature of interactions among cancer cell phenotypes explains the coexistence of diversity in tumors [86, 144] and increases waiting times for neoplasms to develop [145]. An extension of the GLV framework with spatial degrees of freedom would provide valuable insight into the conditions for persistent heterogeneity and how it relates with cancer resilience to perturbations.
9. As it occurs in ecological systems, models of cancer progression often display tipping points separating tumor growth from extinction (or different regimes of growth). What can be learned from the study of tipping points and catastrophic shifts as a way to approach cancer therapies? It has been suggested that we can actually use shifts to “turn ecology against cancer” [146]. Indeed, the potential success of some cancer treatments might be grounded in the possibility of crossing bifurcation points leading to non-viable (or stagnation) states [19, 23]. What is the effect of considering a multispecies scenario? What are the conditions under which a rich cancer cell population will cross a tipping point after a given therapeutic approach?
10. Ecological communities are often seen as the result of an assembly process leading to a directional sequence of transitions. This so called *ecological succession* refers to a process where a set of populations undergoes a series of changes that follow predictable paths after an initial colonization event in a given habitat, which could be an abandoned field. As discussed in [147], this has a clear parallel in cancer, where this initial event would correspond to the establishment of a primary tumor or secondary metastatic tumor. The use of the GLV formalism would be very helpful to address this problem against available single-cell data on growing tumors. Some useful metrics have already been proposed to quantify the directionality (the arrow of time) of complex multispecies communities [99].
11. Over the last decade, a successful approach to ecosystem complexity has emerged from the analysis of synthetic ecologies obtained from sampling actual communities and growing them in cell cultures [42, 148]. By studying the dynamics of these in vitro ecologies, it has been possible to validate several general principles of community ecology using a combination of experiments and GLV models. These experiments have confirmed the mathematical approximations made by canonical models of tipping points, cooperation or extinction dynamics. Could we consider building synthetic cancer communities to perform similar experiments in the test tube? The emerging science of microbiomes [149] (where the GLV approach has been widely adopted) and the possibilities of metagenomic characterization of their complexity could inspire an analogous research within tumor dynamics.

Overall, the present work proposes a necessary bridge between theoretical community ecology and cancer research. Applying the GLV framework to quantitative tumor systems will bring novel understanding and, more importantly, a more nuanced framework to design therapies targeting ecosystem-level properties of cancer.

## Acknowledgements

G.A.-G. thanks M. Barbier, S. Kéfi, V. Maull and J. Piñero for valuable discussions and L. Feinberg for inspiration. G.A.-G. was supported by a 2022 postdoctoral fellowship of the Fundación Ramón Areces. A.R.A. Anderson gratefully acknowledges funding from the NCI via the Cancer Systems Biology Consortium (CSBC) U54CA274507 and support from the Moffitt Center of Excellence for Evolutionary Therapy. R Solé thanks Serguei Saavedra, Jie Deng and Chengyi Long and the members of the Complex Systems Lab for useful insights and discussions, to Michael O’Riordan for inspiration and the support of the Santa Fe Institute.

## References

[1] G. B. Pierce and W. C. Speers, Cancer research 48, 1996 (1988).

[2] L. M. Merlo, J. W. Pepper, B. J. Reid, and C. C. Maley, Nature reviews cancer 6, 924 (2006).

[3] A. R. Anderson and V. Quaranta, Nature Reviews Cancer 8, 227 (2008).

[4] G. Aguadé-Gorgorio, J. Costa, and R. Solé, BioEssays 45, 2200215 (2023).

[5] S. R. Amend and K. J. Pienta, Oncotarget 6, 9669 (2015).

[6] F. R. Adler and D. M. Gordon, Current opinion in systems biology 17, 1 (2019).

[7] K. V. Myers, K. J. Pienta, and S. R. Amend, Cancer Control 27, 1073274820911058 (2020).

[8] R. A. Gatenby, A. S. Silva, R. J. Gillies, and B. R. Frieden, Cancer research 69, 4894 (2009).

[9] P. T. Hamilton, B. R. Anholt, and B. H. Nelson, Nature Reviews Immunology 22, 765 (2022).

[10] S. A. Levin, Ecosystems 1, 431 (1998).

[11] H. Kitano, Nature 426, 125 (2003).

[12] A. Marusyk, V. Almendro, and K. Polyak, Nature reviews cancer 12, 323 (2012).

[13] A. Marusyk, M. Janiszewska, and K. Polyak, Cancer cell 37, 471 (2020).

[14] I. Vitale, E. Shema, S. Loi, and L. Galluzzi, Nature medicine 27, 212 (2021).

[15] T. S. Deisboeck and I. D. Couzin, Bioessays 31, 190 (2009).

[16] C. S. Holling, Annual review of ecology and systematics 4, 1 (1973).

[17] R. A. Gatenby, Cancer Research 51, 2542 (1991).

[18] R. Eftimie, J. L. Bramson, and D. J. Earn, Bulletin of mathematical biology 73, 2 (2011).

[19] R. Solé and G. Aguadé-Gorgorio, Journal of Theoretical Biology 511, 110552 (2021).

[20] E. Kim, J. S. Brown, Z. Eroglu, and A. R. Anderson, Cancers 13, 823 (2021).

[21] R. P. Garay and R. Lefever, Journal of theoretical biology 73, 417 (1978).

[22] V. A. Kuznetsov, I. A. Makalkin, M. A. Taylor, and A. S. Perelson, Bulletin of mathematical biology 56, 295 (1994).

[23] R. V. Solé and T. S. Deisboeck, Journal of Theoretical Biology 228, 47 (2004).

[24] B. P. Kotler and J. S. Brown, Cancer Control 27, 1073274820951776 (2020).

[25] N. D. Lee, K. Kaveh, and I. Bozic, in Seminars in Cancer Biology (Elsevier, 2023).

[26] R. Mathur, Q. Wang, P. G. Schupp, A. Nikolic, S. Hilz, C. Hong, N. R. Grishanina, D. Kwok, N. O. Stevers, Q. Jin, et al., Cell 187, 446 (2024).

[27] J. West, F. Rentzeperis, C. Adam, R. Bravo, K. A. Luddy, M. Robertson-Tessi, and A. R. Anderson, bioRxiv pp. 2022–06 (2022).

[28] R. Axelrod, D. E. Axelrod, and K. J. Pienta, Proceedings of the National Academy of Sciences 103, 13474 (2006).

[29] A. Chapman, L. F. del Ama, J. Ferguson, J. Kamarashev, C. Wellbrock, and A. Hurlstone, Cell reports 8, 688 (2014).

[30] D. Basanta and A. R. Anderson, Interface focus 3, 20130020 (2013).

[31] H. Byrne, T. Alarcon, M. Owen, S. Webb, and P. Maini, Philosophical Transactions of the Royal Society A: Mathematical, Physical and Engineering Sciences 364, 1563 (2006).

[32] Nonlinearity 23, R1 (2009).

[33] H. M. Byrne, Nature Reviews Cancer 10, 221 (2010).

[34] T. S. Deisboeck, Z. Wang, P. Macklin, and V. Cristini, Annual review of biomedical engineering 13, 127 (2011).

[35] D. Basanta, H. Hatzikirou, and A. Deutsch, The European Physical Journal B 63, 393 (2008).

[36] J. M. Pacheco, F. C. Santos, and D. Dingli, Interface focus 4, 20140019 (2014).

[37] M. Archetti and K. J. Pienta, Nature Reviews Cancer 19, 110 (2019).

[38] K. Staňkova, J. S. Brown, W. S. Dalton, and R. A. Gatenby, JAMA oncology 5, 96 (2019).

[39] G. Bunin, Physical Review E 95, 042414 (2017).

[40] M. Barbier, J.-F. Arnoldi, G. Bunin, and M. Loreau, Proceedings of the National Academy of Sciences 115, 2156 (2018).

[41] J. Deng, M. T. Angulo, and S. Saavedra, ISME communications 1, 22 (2021).

[42] J. Hu, D. R. Amor, M. Barbier, G. Bunin, and J. Gore, Science 378, 85 (2022).

[43] C. D. Gatenbee, A.-M. Baker, R. O. Schenck, M. Strobl, J. West, M. P. Neves, S. Y. Hasan, E. Lakatos, P. Martinez, W. C. Cross, et al., Nature Communications 13, 1798 (2022).

[44] A. R. Anderson, M. A. Chaplain, E. L. Newman, R. J. Steele, and A. M. Thompson, Computational and mathematical methods in medicine 2, 129 (2000).

[45] M. A. Leibold, M. Holyoak, N. Mouquet, P. Amarasekare, J. M. Chase, M. F. Hoopes, R. D. Holt, J. B. Shurin, R. Law, D. Tilman, et al., Ecology letters 7, 601 (2004).

[46] G. Aguadé-Gorgorió and R. Solé, Journal of the Royal Society Interface 17, 20200736 (2020).

[47] E. B. Gunnarsson, S. De, K. Leder, and J. Foo, Journal of theoretical biology 490, 110162 (2020).

[48] J. West, M. Robertson-Tessi, and A. R. Anderson, Trends in Cell Biology 33, 300 (2023).

[49] G. Aguadé-Gorgorió and R. Solé, Evolutionary applications 11, 1283 (2018).

[50] R. Durrett and R. Durrett, Branching process models of cancer (Springer, 2015).

[51] S. A. Levin, Ecology 73, 1943 (1992).

[52] R. M. May, Stability and complexity in model ecosystems (Princeton university press, 2019).

[53] T. J. Case, Ecology 80, 2848 (1999).

[54] A. Tabi, F. Pennekamp, F. Altermatt, R. Alther, E. A. Fronhofer, K. Horgan, E. Mächler, M. Pontarp, O. L. Petchey, and S. Saavedra, Nature Ecology & Evolution 4, 1036 (2020).

[55] R. M. May, Nature 238, 413 (1972).

[56] E. H. van Nes and M. Scheffer, The American Naturalist 164, 255 (2004).

[57] C. A. Serván, J. A. Capitan, J. Grilli, K. E. Morrison, and S. Allesina, Nature ecology & evolution 2, 1237 (2018).

[58] S. H. Strogatz, Nonlinear dynamics and chaos: with applications to physics, biology, chemistry, and engineering (CRC press, 2018).

[59] D. A. Kessler and N. M. Shnerb, Physical Review E 91, 042705 (2015).

[60] F. Roy, M. Barbier, G. Biroli, and G. Bunin, PLoS computational biology 16, e1007827 (2020).

[61] E. Mallmin, A. Traulsen, and S. De Monte, arXiv preprint arXiv:2306.11031 (2023).

[62] I. A. Rodriguez-Brenes, N. L. Komarova, and D. Wodarz, Trends in ecology & evolution 28, 597 (2013).

[63] Z. Frankenstein, D. Basanta, O. E. Franco, Y. Gao, R. A. Javier, D. W. Strand, M. Lee, S. W. Hayward, G. Ayala, and A. R. Anderson, Nature Ecology & Evolution 4, 870 (2020).

[64] B. Jansson and L. Révész, Mathematical Biosciences 19, 131 (1974).

[65] Ž. Bajzer, S. Vuk-Pavlović, and M. Huzak, Mathematical modeling of tumor growth kinetics (1997).

[66] L. De Pillis, Bulletin of mathematical biology 76, 2010 (2014).

[67] R. A. Gatenby, Journal of theoretical biology 176, 447 (1995).

[68] R. Gatenby, European Journal of Cancer 32, 722 (1996).

[69] J. A. Spratt, D. Von Fournier, J. S. Spratt, and E. E. Weber, Cancer 71, 2013 (1993).

[70] P. Gerlee and A. R. Anderson, Physical biology 12, 056001 (2015).

[71] I. Scheuring and E. Szathmary, Journal of theoretical biology 212, 99 (2001).

[72] E. Szathmáry and J. M. Smith, Journal of theoretical biology 187, 555 (1997).

[73] I. A. Hatton, O. Mazzarisi, A. Altieri, and M. Smerlak, Science 383, eadg8488 (2024).

[74] T. B. Taylor, A. V. Wass, L. J. Johnson, and P. Dash, BMC evolutionary biology 17, 1 (2017).

[75] B. J. Hershey, S. Barozzi, F. Orsenigo, S. Pompei, F. Iannelli, S. Kamrad, V. Matafora, F. Pisati, L. Calabrese, G. Fragale, et al., Science Advances 9, eadh4184 (2023).

[76] J. Salimi Sartakhti, M. H. Manshaei, D. Basanta, and M. Sadeghi, PloS one 12, e0175063 (2017).

[77] H. Cho, A. L. Lewis, K. M. Storey, and H. M. Byrne, Journal of Theoretical Biology 559, 111377 (2023).

[78] H. Tari, K. Kessler, N. Trahearn, B. Werner, M. Vinci, C. Jones, and A. Sottoriva, Cell Reports 40 (2022).

[79] R. M. May, Science 241, 1441 (1988).

[80] R. Margalef, Oldendorf: Ecology Institute. (1997).

[81] K. S. McCann, Nature 405, 228 (2000).

[82] R. A. Armstrong and R. McGehee, The American Naturalist 115, 151 (1980).

[83] R. MacArthur and R. Levins, Proceedings of the National Academy of Sciences 51, 1207 (1964).

[84] A. Marusyk, D. P. Tabassum, P. M. Altrock, V. Almendro, F. Michor, and K. Polyak, Nature 514, 54 (2014).

[85] A. S. Cleary, T. L. Leonard, S. A. Gestl, and E. J. Gunther, Nature 508, 113 (2014).

[86] I. González-García, R. V. Solé, and J. Costa, Proceedings of the National Academy of Sciences 99, 13085 (2002).

[87] N. McGranahan and C. Swanton, Cell 168, 613 (2017).

[88] D. Wodarz and N. Komarova, Dynamics of cancer: mathematical foundations of oncology (World Scientific, 2014).

[89] E. Kim, V. Rebecca, I. V. Fedorenko, J. L. Messina, R. Mathew, S. S. Maria-Engler, D. Basanta, K. S. Smalley, and A. R. Anderson, Cancer research 73, 6874 (2013).

[90] A. E. Magurran and P. A. Henderson, Nature 422, 714 (2003).

[91] J. Gallaher, M. Strobl, J. West, J. Zhang, R. Gatenby, M. Robertson-Tessi, and A. R. Anderson, bioRxiv pp. 2022–08 (2022).

[92] L. J. Gay and I. Malanchi, Biochimica et Biophysica Acta (BBA)-Reviews on Cancer 1868, 231 (2017).

[93] P. Sharma, S. Hu-Lieskovan, J. A. Wargo, and A. Ribas, Cell 168, 707 (2017).

[94] R. J. Gillies and R. A. Gatenby, Journal of bioenergetics and biomembranes 39, 251 (2007).

[95] T. Epstein, R. A. Gatenby, and J. S. Brown, PloS one 12, e0185085 (2017).

[96] W. Lopes, D. Amor, and J. Gore, bioRxiv pp. 2023–12 (2023).

[97] M. J. Williams, B. Werner, C. P. Barnes, T. A. Graham, and A. Sottoriva, Nature genetics 48, 238 (2016).

[98] P. B. Gupta, I. Pastushenko, A. Skibinski, C. Blanpain, and C. Kuperwasser, Cell stem cell 24, 65 (2019).

[99] G. Bunin, Oikos 130, 489 (2021).

[100] J. Huisman and F. J. Weissing, The American Naturalist 157, 488 (2001).

[101] C. C. Chang and B. L. Turner, Ecological succession in a changing world (2019).

[102] C. Song, S. Von Ahn, R. P. Rohr, and S. Saavedra, Trends in Ecology & Evolution 35, 384 (2020).

[103] C. Long, J. Deng, J. Nguyen, Y.-Y. Liu, E. J. Alm, R. Solé, and S. Saavedra, Proceedings of the National Academy of Sciences 121, e2312521121 (2024).

[104] C. Aktipis, A. M. Boddy, R. A. Gatenby, J. S. Brown, and C. C. Maley, Nature Reviews Cancer 13, 883 (2013).

[105] R. V. Solé, J. M. Montoya, and D. H. Erwin, Philosophical Transactions of the Royal Society of London. Series B: Biological Sciences 357, 697 (2002).

[106] K. Liautaud, E. H. van Nes, M. Barbier, M. Scheffer, and M. Loreau, Ecology letters 22, 1243 (2019).

[107] M. Strobl, J. Gallaher, M. Robertson-Tessi, J. West, and A. Anderson, Annals of Oncology 34, 867 (2023).

[108] J. Househam, T. Heide, G. D. Cresswell, I. Spiteri, C. Kimberley, L. Zapata, C. Lynn, C. James, M. Mossner, J. Fernandez-Mateos, et al., Nature 611, 744 (2022).

[109] C. Neftel, J. Laffy, M. G. Filbin, T. Hara, M. E. Shore, G. J. Rahme, A. R. Richman, D. Silverbush, M. L. Shaw, C. M. Hebert, et al., Cell 178, 835 (2019).

[110] N. Q. Balaban, J. Merrin, R. Chait, L. Kowalik, and S. Leibler, Science 305, 1622 (2004).

[111] C. Scheel and R. A. Weinberg, International journal of cancer 129, 2310 (2011).

[112] S. V. Sharma, D. Y. Lee, B. Li, M. P. Quinlan, F. Takahashi, S. Maheswaran, U. McDermott, N. Azizian, L. Zou, M. A. Fischbach, et al., Cell 141, 69 (2010).

[113] A. Goldman, B. Majumder, A. Dhawan, S. Ravi, D. Goldman, M. Kohandel, P. K. Majumder, and S. Sengupta, Nature communications 6, 6139 (2015).

[114] D. B. Burkhardt, B. P. San Juan, J. G. Lock, S. Krishnaswamy, and C. L. Chaffer, Cancer discovery 12, 1847 (2022).

[115] G. Aguadé-Gorgorio, S. Kauffman, and R. Solé, Bulletin of mathematical biology 84, 24 (2022).

[116] I. Smalley, E. Kim, J. Li, P. Spence, C. J. Wyatt, Z. Eroglu, V. K. Sondak, J. L. Messina, N. A. Babacan, S. S. Maria-Engler, et al., EBioMedicine 48, 178 (2019).

[117] A. S. Perelson and G. Weisbuch, Reviews of modern physics 69, 1219 (1997).

[118] Y. Yang et al., The Journal of clinical investigation 125, 3335 (2015).

[119] T. N. Schumacher and R. D. Schreiber, Science 348, 69 (2015).

[120] L. Zapata, G. Caravagna, M. J. Williams, E. Lakatos, K. AbdulJabbar, B. Werner, D. Chowell, C. James, L. Gourmet, S. Milite, et al., Nature Genetics 55, 451 (2023).

[121] N. McGranahan, A. J. Furness, R. Rosenthal, S. Ramskov, R. Lyngaa, S. K. Saini, M. Jamal-Hanjani, G. A. Wilson, N. J. Birkbak, C. T. Hiley, et al., Science 351, 1463 (2016).

[122] E. Lakatos, M. J. Williams, R. O. Schenck, W. C. Cross, J. Househam, L. Zapata, B. Werner, C. Gatenbee, M. Robertson-Tessi, C. P. Barnes, et al., Nature Genetics 52, 1057 (2020).

[123] D. S. Chen and I. Mellman, Nature 541, 321 (2017).

[124] M. A. Nowak, R. M. Anderson, A. R. McLean, T. F. Wolfs, J. Goudsmit, and R. M. May, Science 254, 963 (1991).

[125] D. Basanta, D. W. Strand, R. B. Lukner, O. E. Franco, D. E. Cliffel, G. E. Ayala, S. W. Hayward, and A. R. Anderson, Cancer research 69, 7111 (2009).

[126] P. D. Katsikis, K. J. Ishii, and C. Schliehe, Nature Reviews Immunology 24, 213 (2024).

[127] S. R. Amend, S. Roy, J. S. Brown, and K. J. Pienta, Cancer letters 380, 237 (2016).

[128] A. Heyde, J. G. Reiter, K. Naxerova, and M. A. Nowak, Proceedings of the National Academy of Sciences 116, 14129 (2019).

[129] I. Hanski, Metapopulation ecology (Oxford University Press, 1999).

[130] M. Barbier, G. Bunin, and M. A. Leibold, bioRxiv pp. 2023–06 (2023).

[131] A. Marusyk and K. Polyak, Biochimica et Biophysica Acta (BBA)-Reviews on Cancer 1805, 105 (2010).

[132] G. G. Lorenzana, A. Altieri, and G. Biroli, arXiv preprint arXiv:2309.09900 (2023).

[133] N. S. Goel, S. C. Maitra, and E. W. Montroll, Reviews of modern physics 43, 231 (1971).

[134] G. Biroli, G. Bunin, and C. Cammarota, New Journal of Physics 20, 083051 (2018).

[135] R. Lande, S. Engen, and B.-E. Saether, Stochastic population dynamics in ecology and conservation (Oxford University Press, USA, 2003).

[136] R. V. Solé, D. Alonso, and A. McKane, Philosophical Transactions of the Royal Society of London. Series B: Biological Sciences 357, 667 (2002).

[137] A. Hastings and K. Higgins, Science 263, 1133 (1994).

[138] R. V. Solé, J. Bascompte, and J. Valls, Chaos: An Interdisciplinary Journal of Nonlinear Science 2, 387 (1992).

[139] L. A. Saravia, G. D. Ruxton, and C. E. Coviella, Proceedings of the Royal Society of London. Series B: Biological Sciences 267, 1781 (2000).

[140] T. B. Francis, K. C. Abbott, K. Cuddington, G. Gellner, A. Hastings, Y.-C. Lai, A. Morozov, S. Petrovskii, and M. L. Zeeman, Nature Ecology & Evolution 5, 285 (2021).

[141] D. Tilman and P. Kareiva, Spatial ecology: the role of space in population dynamics and interspecific interactions (Princeton University Press, 1997).

[142] J. Bascompte and R. Solé, Modeling spatiotemporal dynamics in ecology (1998).

[143] R. V. Solé, J. Bascompte, and J. Valls, Journal of theoretical Biology 159, 469 (1992).

[144] B. Waclaw, I. Bozic, M. E. Pittman, R. H. Hruban, B. Vogelstein, and M. A. Nowak, Nature 525, 261 (2015).

[145] E. A. Martens, R. Kostadinov, C. C. Maley, and O. Hallatschek, New journal of physics 13, 115014 (2011).

[146] K. S. Korolev, J. B. Xavier, and J. Gore, Nature Reviews Cancer 14, 371 (2014).

[147] I. Kareva, Biological Theory 10, 283 (2015).

[148] T. Großkopf and O. S. Soyer, Current opinion in microbiology 18, 72 (2014).

[149] S. Widder, R. J. Allen, T. Pfeiffer, T. P. Curtis, C. Wiuf, W. T. Sloan, O. X. Cordero, S. P. Brown, B. Momeni, W. Shou, et al., The ISME journal 10, 2557 (2016).

